# An optimized competitive-aging method reveals gene-drug interactions underlying the chronological lifespan of *Saccharomyces cerevisiae*

**DOI:** 10.1101/696682

**Authors:** J. Abraham Avelar-Rivas, Michelle Munguía-Figueroa, Alejandro Juárez-Reyes, Erika Garay, Sergio E. Campos, Noam Shoresh, Alexander DeLuna

## Abstract

The chronological lifespan of budding yeast is a model of aging and age-related diseases. This paradigm has recently allowed genome-wide screening of genetic factors underlying post-mitotic viability in a simple unicellular system, which underscores its potential to provide a comprehensive view of the aging process. However, results from different large-scale studies show little overlap and typically lack quantitative resolution to derive interactions among different aging factors. We previously introduced a sensitive, parallelizable approach to measure the chronological-lifespan effects of gene deletions based on the competitive aging of fluorescence-labeled strains. Here, we present a thorough description of the method, including an improved multiple-regression model to estimate the association between death rates and fluorescent signals, which accounts for possible differences in growth rate and experimental batch effects. We illustrate the experimental procedure—from data acquisition to calculation of relative survivorship—for ten deletion strains with known lifespan phenotypes, which is achieved with high technical replicability. We apply our method to screen for gene-drug interactions in an array of yeast deletion strains, which reveals a functional link between protein glycosylation and lifespan extension by metformin. Competitive-aging screening coupled to multiple-regression modeling provides a powerful, straight-forward way to identify aging factors in yeast and their interactions with pharmacological interventions.

## INTRODUCTION

A major challenge in aging research is to describe the way in which different genetic pathways and biochemical processes mediating aging are interconnected to one another (Kirkwood, 2008; López-Otín et al., 2013). Simple cellular models provide a starting point to grant a systems-level understanding of aging, in which the lifespan phenotype is addressed as a complex trait resulting from the action of multiple genes, cellular processes, environmental factors, and their interactions.

The chronological lifespan (CLS) of *Saccharomyces cerevisiae* is used to describe genetic, nutrimental, and pharmacological factors underlying survivorship of post-mitotic, non-dividing cells (Longo and Fabrizio, 2012). The budding yeast’s replicative-lifespan and CLS are simple experimental models that have been used to reveal the conserved lifespan-extending effects of reduced TOR and RAS/PKA signaling, as well as the anti-aging effect of rapamycin, spermidine, and caloric restriction (Wei et al., 2008; Eisenberg et al., 2009; Longo and Fabrizio, 2012; Gems and Partridge, 2013). Traditionally, the CLS of a yeast-cell population is measured by counting colony-forming units from samples of a long-term stationary-phase culture (Longo et al., 2012; Hu et al., 2013). More recently, large-scale screening approaches have been implemented to screen for genetic aging factors in yeast. These studies provide unbiased catalogues of CLS mutant phenotypes (Powers et al., 2006; Fabrizio et al., 2010; Matecic et al., 2010; Burtner et al., 2011; Garay et al., 2014), mutants with diminished or enhanced response to dietary restriction or nutrient limitation (Gresham et al., 2011; Campos et al., 2018), and CLS phenotypes of collections of wild isolates and lines derived from biparental crosses (Jung et al., 2018; Barré et al., 2019).

A current limitation in the field is that large-scale CLS-phenotyping screens have resulted in a large number of false positive hits when further confirmed by smaller-scale approaches, ranging from 50% to 94% (Powers et al., 2006; Fabrizio et al., 2010; Matecic et al., 2010; Burtner et al., 2011; Garay et al., 2014). In addition, comparisons of different large-scale studies show that there is little overlap among the identified genetic factors, which could be explained in part by differences in genotypic background, media composition, and subtle environmental variations (Smith et al., 2016). In addition, changes in controlled or uncontrolled environmental conditions are known to be important modifiers of CLS phenotypes and confounding causes of aging (Burtner et al., 2009, 2011; Santos et al., 2015; Smith et al., 2016; Campos and DeLuna, 2019). In this context, a combination of high throughput and resolution is much needed to correctly determine not only genetic aging factors, but also to quantitatively derive their interactions with nutrimental, chemical, or pharmacological environments.

In an effort to improve the throughput of CLS screening without sacrificing phenotyping sensitivity, we previously introduced a competition-based method for quantitative large-scale genetic analysis (Garay et al., 2014). In brief, each RFP-labeled single-deletion strain is mixed with a CFP wild-type reference and grown to saturation; fluorescence signal in outgrowth cultures is used to estimate the relative number of viable cells in the non-dividing culture at different time points in stationary phase. This approach recapitulates known CLS factors and suggests new lifespan phenotypes in yeast (Garay et al., 2014). More recently, we have used this approach to screen for dietary restriction factors, namely CLS gene-deletion phenotypes that are aggravated or alleviated when yeast populations are aged under a poor nitrogen source (Campos et al., 2018).

In this study, we describe an optimized multiple regression modeling strategy to analyze measurements from our competition-based approach for CLS genetic analysis in yeast, by accounting for possible differences in growth rate and experimental batch effects. In addition, we provide a systematic analysis of the method’s replicability and data-analysis scripts. For ten knockout strains, we compare the replicability of our results with those obtained with a useful parallelizable approach based on outgrowth kinetics (Murakami et al., 2008; Jung et al., 2015). Importantly, we take advantage of our improved data-analysis method to derive gene-drug interactions by measuring the relative effects on survival of metformin in 76 deletion strains of widely conserved genes. We discuss the potential of competitive-aging screening to describe large numbers of genetic and environmental interactions underlying aging and longevity in aging cells.

## MATERIALS AND METHODS

### Strains and media

Ten single-gene deletion strains targeting *ATG1*, *HAP3*, *MSN2*, *MSN4*, *RAS2*, *RIM15*, *RPS16A*, *STE12*, *GLN3*, and *SWR1* were generated *de novo* by PCR-based gene replacement in the YEG01-RFP background(Garay et al., 2014) using the *natMX4* module from pAG25 (Euroscarf). In addition, two isogenic reference strains were generated over the YEG01-RFP and YEG01-CFP backgrounds by deleting the neutral *HO* locus. Lithium-acetate transformation and 100 µg/ml clonNAT (Werner BioAgents) were used. The resulting strains were *MAT-*α *x*Δ*::natMX4 PDC1-RFP.CaURA3MX4 can1*Δ*::STE2pr-SpHIS5 lyp1*Δ *ura3Δ0 his3Δ1 LEU2 MET15*; strong constitutive expression of fluorescent proteins during exponential growth is achieved by carboxyl-terminal fusion to the Pdc1 protein.

For gene-drug interactions, a collection of RFP-tagged gene-deletion strains was generated by mating an array of 85 strains from the yeast deletion collection to the *ho*Δ YEG01-RFP SGA-starter strain, as previously described(Garay et al., 2014). Resulting prototrophic strains were *MAT-*a *x*Δ*::kanMX4 PDC1-RFP.CaURA3MX4 can1*Δ*::STE2pr-SpHIS5 lyp1*Δ *ura3Δ0 his3Δ1 LEU2 MET15*. The CFP reference strain was the neutral marker reference *his3*Δ::*kanMX4*. Double-marker strains and CLS data were successfully obtained for 76 deletion strains (Supplementary Data S1).

Aging medium was synthetic complete (SC) with ammonium, 2% glucose, and 0.2% amino acid supplement mix (CSHL Manual, 2005), without buffering. 40mM metformin (Sigma D150959) was used where indicated. Single-culture and competitive outgrowth kinetics were done in low fluorescence YNB medium (YNB-lf) with 2% glucose and complete amino-acids supplement mix (Sigma Y1501 completed with Sigma U0750)(Sheff and Thorn, 2004). All cultures were incubated in a growth chamber at 30°C and 80%-90% relative humidity without shaking; aging co-cultures were vigorously shaken at every sampling point. For media recipes, see Supplementary Table S1.

### Competitive-aging culture setup and outgrowth measurements

To obtain relative CLS measurements, mutant (*x*Δ_RFP_) and reference strains (WT_RFP_ and WT_CFP_) were pre-cultured separately until saturation. Saturated cultures were mixed in a 2:1 mutant:reference ratio and ∼1.5 μL aliquots were transferred with a 96 solid-pin replicator (V&P Scientific VP 407) onto 96 semi deepwell plates (Nunc 260251) with 750 µL of fresh aging medium and disposable plastic covers.

Outgrowth sampling of the competitive-aging culture began 3 days after initial inoculation (time zero) and was repeated initially every 24 h and 48-72 h afterwards for up to 16 days after time zero. At each sampling day, *T_i_*, 12 µL of shaken aging cultures were inoculated onto 140µL fresh YNB-lf medium in clear polystyrene 96-well plates (Corning 3585). Outgrowth was monitored every 1-3 h until cultures reached stationary phase, by measuring raw fluoresence of mCherry (*RFP*, Ex 587/5 nm and Em 610/5 nm), Cerulean (*CFP*, Ex 433/5 nm and Em 475/5 nm), and absorbance at 600 nm (*OD_600_*) using a Tecan Infinite M1000 reader integrated to a robotic platform (Tecan Freedom EVO200). Outgrowth cultures were resuspended by vigorous shaking right before every measurement. To increase fluorescence-signal dynamic range, an outgrowth culture with the samples was measured one day before the actual experiment to calculate the optimal-gain at late exponential-growth, when fluorescence signal is at its maximum level. Optimal gain values were fixed for all measurements in the experiment; values used were 140-167 and 131-137 for *RFP* and *CFP*, respectively.

### Death-rate calculation from competitive-aging outgrowth data

The 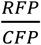 signal ratio was used to estimate the number of cells expressing each fluorescent signal. Auto fluorescence background was defined as the *RFP* and *CFP* signal of WT_CFP_ and WT_RFP_ monocultures, respectively, and was subtracted from all competitive aging cultures. Using data from different measurements in the outgrowth cultures (*t_j_*, in hours) of each stationary-phase sampling point (*T_i_*, in days), the signal ratio 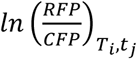 of each sample (*w*) was fitted to the linear model *A_w_ + S_w_· T_i_ + G_w_ ·t_j_ + C_Ti, tj_*. The ratio of viable cells at the beginning of the experiment is modeled in *A*, while *S* (relative survivorship) is the death-rate difference of the mutant and wild-type reference, and *G* (relative growth) is their growth-rate difference in the outgrowth co-culture. In addition, the term *C_Ti,tj_* was introduced to consider the systematic variation of each plate at each stationary-phase sampling point *T_i_t_j_*. A complete description of the model and its implementation is provided in Supplementary Note S1.

### Deriving gene-drug interactions from CLS phenotypes

To identify gene-drug interactions, namely cases in which the CLS phenotype of a gene-deletion strain is significantly aggravated or alleviated by treatment with a drug, the relative survivorship of a set of 76 deletion strains was measured with and without 40 mM metformin. Deletion strains were randomly selected from the yeast deletion collection, considering only genes with a mammalian ortholog (Ensembl). Relative survivorship values were rescaled to a dimensionless parameter, given that metformin increases yeast CLS and that the relative survivorship *S* expressed in days is constrained by the death rate of the wild-type reference, *r_wt_*, which is different in each condition. For each mutant under a given condition, rescaled survivorship (*rS*) was defined as 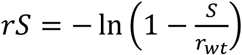, where *r_wt_* is the average death rate of all control wild-type competitions in each plate. Gene-drug interactions were defined as cases in which a mutant’s *rS* in SC was significantly different to the *rS* in SC + metformin (*p*<0.05; *t*-test).

### Measuring death rate in monoculture from outgrowth kinetics

CLS estimates based on outgrowth kinetics of single-population aging cultures were adapted from a well established high-throughput method(Murakami et al., 2008; Jung et al., 2015). Culture density was monitored by measuring absorbance at *OD_600nm_*; background signal—*OD_600_* at outgrowth inoculation—was subtracted to each data point. At each aging sampling point *T_i_* (days), the time-shift in hours, t, to reach a fixed cell density of *OD_600_*=0.35 reports for the remaining fraction of viable cells in the population, as previously reported(Murakami et al., 2008). Specifically, for each successive age time point, the percent of viability (*V_T_*) was calculated using the equation 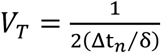, where Δt*_n_* is the time shift and δ is the doubling time. Death rates were the exponential decay rates calculated by fitting all *V_T_* data points as a function of time in stationary phase to a first-order exponential model (Matlab, *fit*).

### Live/dead cell viability assays

Stationary-phase cultures (5 µL) were transferred to 50 µL of BD FACSFlow (BD #342003) with 0.1% propidium iodide (PI) and 0.1% SYTO9 green (ThermoFisher L34952) in 96-well plates (Corning 3585), shaken for 1 min at 1,000 rpm in a plate shaker (Heidolph Titramax 1000), and incubated at room temperature in the dark for 20 minutes. An analytical flow cytometer with a high-throughput sampler (LSRFortessa-HTS, Becton Dickinson) was used to capture 10,000 events. SYTO9 fluorescence was excited with a 488nm blue laser and collected through a 525/50 nm band-pass filter and a 505LP emission filter, while PI signal was excited with a 488nm blue laser and collected through a 712/21 band-pass filter. Events with a increased PI/SYTO9 signal were counted as alive.

## RESULTS

### Relative survivorship in stationary phase can be estimated from bulk fluorescence signal of two populations in co-culture

We sought to optimize data analysis and systematically test a competition-based method aimed at describing CLS phenotypes of viable gene-deletion strains in budding yeast. To directly measure the lifespan effects, we tracked changes in the relative abundances of RFP-deletion (*x*Δ_RFP_) and CFP-tagged wild-type (WT_CFP_) strains in co-culture, as a function of time in stationary phase (*T*, days) (**Figure 1A**). To this end, we inoculated stationary-phase cells at different time points into fresh medium and monitored the outgrowth at multiple times (*t*, hours) by measuring absorbance at 600nm (*OD_600_*), bulk RFP signal (*RFP*), and bulk CFP signal (*CFP*), until the outgrowth co-cultures reached saturation.

**Figure 1.**
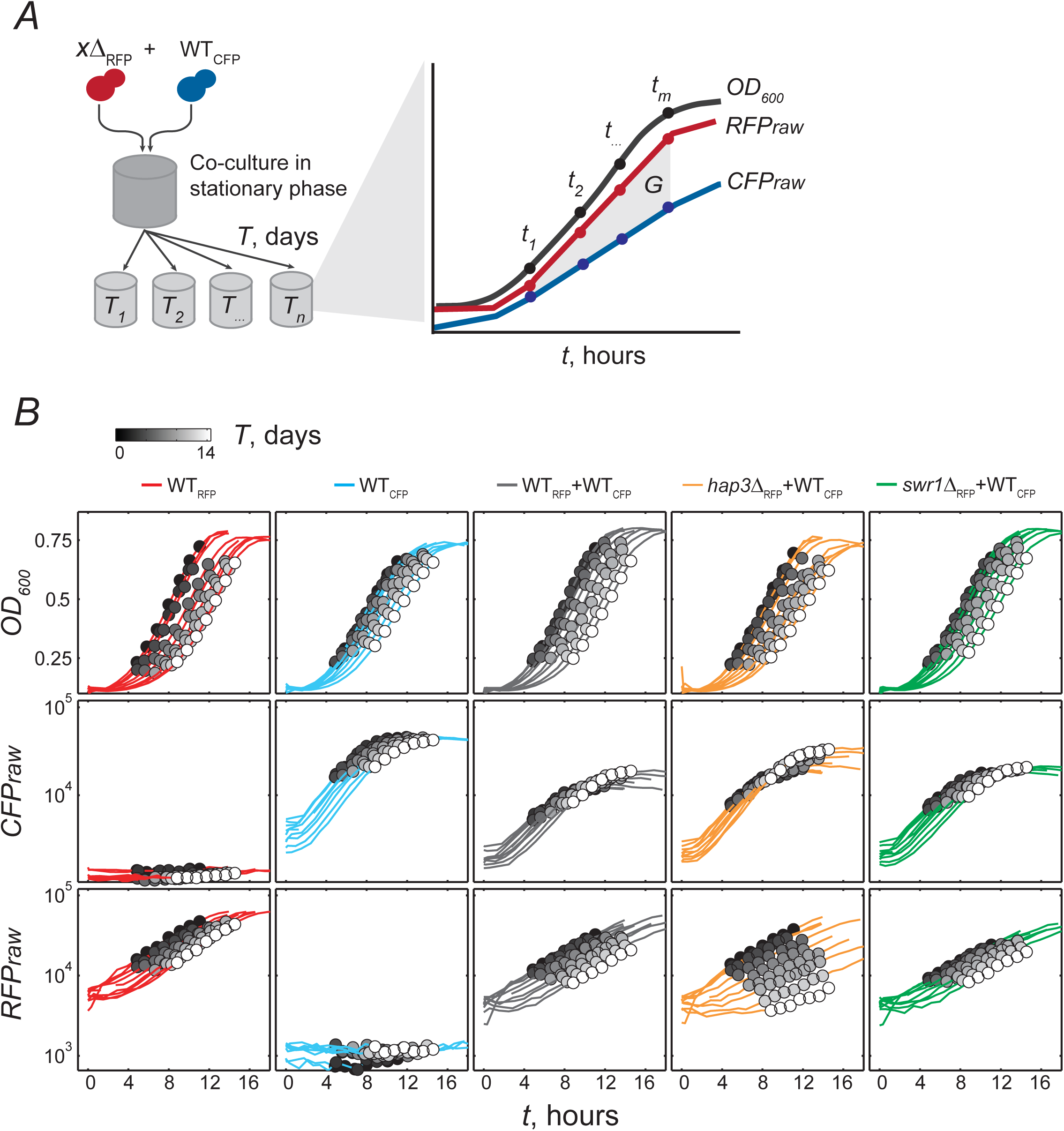
Competitive-aging of fluorescently labeled budding yeast strains. ***A***, Schematic of the experimental setup. A co-culture of fluorescence-tagged gene deletion (*x*Δ_RFP_) and wild-type reference (WT*_CFP_*) strains in stationary phase is sampled regularly at days *T*. Outgrowth cultures in fresh medium are monitored at *t* hours with simultaneous measurement of absorbance at 600nm (*OD_600_*), and raw *RFP* and *CFP* signals. Change in fluorescent-signal ratio in the outgrowth cultures is used to estimate the relative number of viable cells after *T* days in stationary phase. Possible differences in growth rate (*G*) are taken into account (see multiple linear modeling). ***B,*** Raw data of *OD_600_*, *RFPraw*, and *CFPraw* signal obtained from outgrowth-culture kinetics started after *T* days in stationary phase. Selected examples are shown in five columns, from left to right: Only WT_RFP_ (red lines), only WT_CFP_ (blue lines), both WT populations in co-culture (WT_RFP_ +WT_CFP_, gray lines), a short-lived deletion strain in co-culture with the WT (*hap3*Δ_RFP_ +WT_CFP_, orange lines), and a long-lived strain with the WT (*swr1*Δ_RFP_ +WT_CFP_, green lines). Samples are measured throughout the outgrowth culture after inoculation, but analysis only considers data points at exponential growth (circles in grayscale).

We characterized the CLS of ten deletion strains that are known to show increased or decreased lifespan (Fabrizio and Longo, 2003; Wei et al., 2008; Alvers et al., 2009; Laschober et al., 2010; Burtner et al., 2011; Garay et al., 2014; Campos et al., 2018). Each *x*Δ_RFP_ was co-cultured until saturation with the WT_CFP_ reference in up to seven replicates in a single deep-well plate (see Materials and Methods). Competitive-aging cultures were monitored for ∼15 days in stationary phase. As expected, outgrowth kinetics measured by *OD_600_* showed a clear shift with time (days) in stationary phase; aging co-cultures gradually took a longer time in outgrowth (hours) to reach a given cell density (**Figure 1B**). This prevalent shift in growth kinetics reflects the loss of viability with culture age, as previously described (Murakami et al., 2008).

In terms of fluorescence-signal kinetics, we observed that the WT_RFP_ or WT_CFP_ monocultures mostly recapitulated *OD_600_* kinetics, suggesting that loss of viability can also be measured by the shift of the fluorescence signal over the days (**Figure 1B**, red and blue lines). WT_RFP_+WT_CFP_ populations competed in co-culture showed similar shifts in fluorescence signals, suggesting that loss of viability occurred at similar rates in both wild-type populations, as expected (**Figure 1B**, gray lines). In contrast, some *x*Δ_RFP_+WT_CFP_ co-cultures showed delayed or exacerbated shifts in fluorescence kinetics. For instance, the *hap3*Δ_RFP_+WT_CFP_ co-culture showed a steep decrease in bulk RFP signal as a function of days in stationary phase, along with a slight increase in CFP signal (**Figure 1B**, orange lines), suggesting that loss of viability occurred at a rate faster than the WT_CFP_ reference (short-lifespan phenotype). In contrast, the *swr1*Δ_RFP_+WT_CFP_ co-culture showed a small increase in RFP signal along with steady CFP signal (**Figure 1B**, green lines), suggesting slower loss of viability of *swr1*Δ compared to the WT (long-lifespan phenotype). These examples illustrate, from raw data, that changes in bulk fluorescence in co-culture can be used to estimate loss of cell population viability as a function of time in stationary phase.

### A model to adjust relative survivorship in co-culture from changes in relative fluorescence signal

To obtain a quantitative phenotypic value from our experimental measurements, we developed a model that provides a relative survivorship parameter (*S*). Specifically, we established a multiple linear regression analysis where each experimental measurement in the outgrowth culture is modeled as:

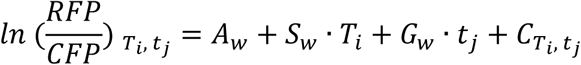

Using a system of linear equations, we obtained the regression coefficients using all outgrowth measurements in a 96-well plate. The contribution of relative survivorship (*S*) along with that of the other three parameters for competing populations are illustrated in **Figure 2**. The expected value, 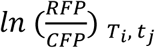, is the logarithmic quotient of the sizes of the populations of *x*Δ_RFP_ and WT_CFP_ from an outgrowth inoculated at day *T_i_* in stationary phase and measured after *t_j_* hours in the outgrowth culture (Supplementary Note S1). The parameter *S* is the difference in death rates (**Figure 2A**, linear regression, open circles), while *G* is the difference in growth rates (linear regression, crosses in **Fig2A,B**). The parameter *A_w_* is the logarithmic proportion of the sizes of both populations at the beginning of the experiment (**Figure 2B**, solid vertical lines at *T*_1_ = 0). The term *C_Ti,tj_* is the error in each measurement, which is mostly determined by the deviation from zero change in WT_RFP_+WT_CFP_ reference competitions, and is similar to the deviation of measurements in *x*Δ_RFP_+WT_CFP_ competitions (**Figure 2B**). The rationale here is that relative survivorship (*S*) and relative growth rate (*S*) are by definition equal to zero in reference competitions (WT_RFP_+WT_CFP_), and therefore consistent changes in 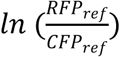 are likely due to systematic errors in the measurements. It must be noted that there are usually survivorship and growth-rate differences between the two wild-type strains (WT_RFP_ and WT_CFP_), which are taken into account while fitting the data. For instance, in this experiment the WT_RFP_ (and isogenic deletion strains) die faster than the WT_CFP_ reference used, which is evident from the negative slope of data points in **Fig 2A**; this lifespan difference is corrected by fitting data to the model in which the WT has *S* =0.

**Figure 2.**
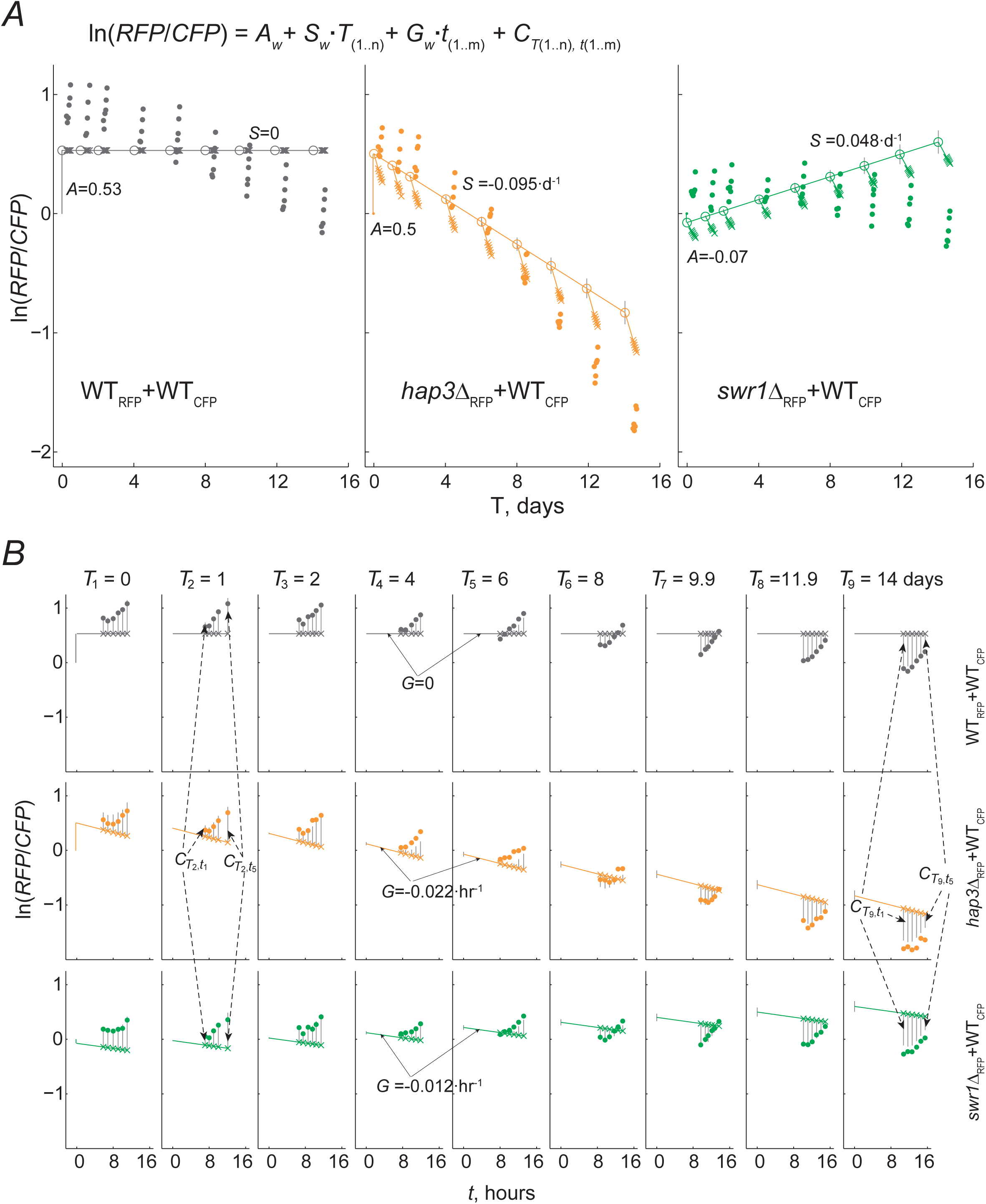
Multiple linear modeling fits relative survivorship and relative growth rate from competitive-aging cultures. Examples of a reference competition (gray) and two deletion strains in co-culture with the WT_CFP_ (*hap3*Δ_RFP_, orange; *swr1*Δ_RFP_, green). In both panels, dots indicate the experimental *ln*(*RFP/CFP*) measurements. ***A*,** Fitted data is shown in the *T* timescale (days). Open circles are the data fitted at *T_i_* and error bars at those time points are the 95% CI of the fit; their linear regression has slope *S* (the relative survivorship, our parameter of interest). For each well, parameter *A* indicates the deviation from ln(*RFP/CFP*)=0 at *T* =0, as indicated by the vertical line at *T* =0. ***B*,** Fitted data is shown in the *t* timescale (hours); nine panels correspond to outgrowths at different age times, *T*. Crosses shown directly below or above experimental data are the fitted data 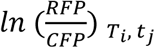 at the time of each measurement; their linear regression has slope *G* (the relative growth rate). All samples (wells) in the same 96-well plate measurement have the same *C_T,t_* (systematic batch error, gray vertical lines).

In the following sections, we show that competitive-aging experiments with multiple-regression modeling provide reliable and replicable quantifications of relative survivorship. We also show that this procedure is useful to identify CLS phenotypes and to score their interactions with pharmacological factors.

### Competitive-aging experiments provide replicable estimates of survivorship

To systematically assess the technical replicability of the competitive-aging method, we measured the CLS of ten mutants and reference strains in 96-well plates with multiple independent replicate wells. Specifically, we measured relative survivorship of up to seven replicate samples of each one of the ten deletion mutants together with 31 wild-type reference competitions in a 96-well plate (**Figure 3A**); the entire experiment was carried two times: one experiment with three replicate plates and the second one with two replicate plates. We validated our results with another large-scale method that provides precise estimates of survivorship (**Figure 3B**). In particular, we measured CLS of the same array of mutants and wild-type strains using an established high-throughput method that is based in the changes of outgrowth kinetics of aging monocultures (Murakami et al., 2008; Jung et al., 2015). Qualitative inspection showed that both methods correctly scored those mutants with known CLS effect (Fabrizio et al., 2001; Fabrizio and Longo, 2003). Specifically *rim15*Δ and *hap3*Δ had reduced survivorship, while *ras2*Δ and *gln3*Δ showed increased lifespan compared to wild type.

**Figure 3.**
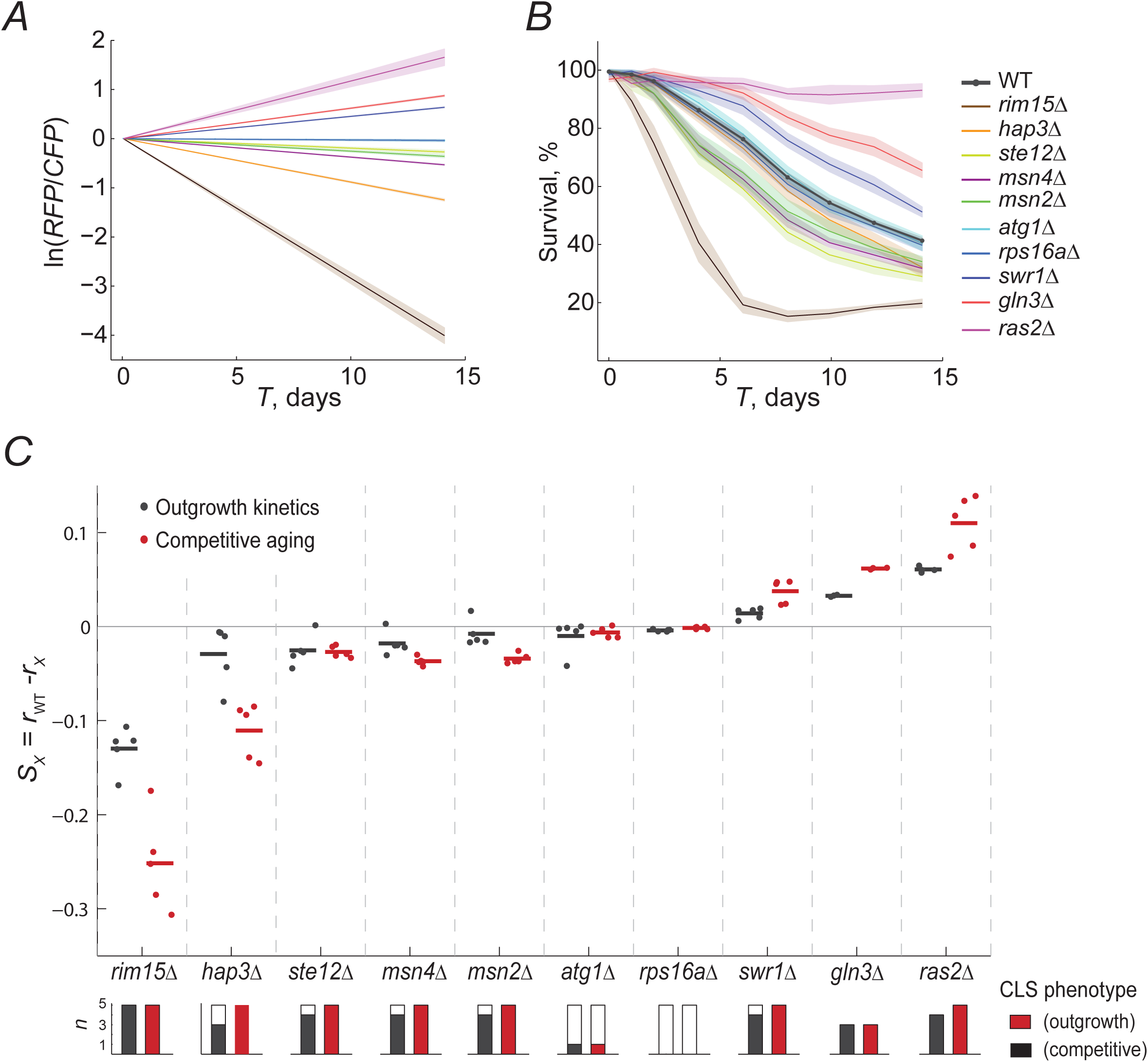
The competitive-aging method is accurate and replicable. ***A***, Relative survivorship estimated by competitive aging. The modeled *ln*(*RFP*/*CFP*) is shown for the WT and ten mutant strains at each T_i_ point, averaged for at least six technical replicates along with the CI 95% (shaded area). ***B***, Percent of surviving cells in monoculture over time measured by the shift in outgrowth kinetics; the mean and CI 95% are shown. ***C***, Average relative survivorship of three to five experimental replicates is shown, expressed either as the difference of individual death rates for monoculture outgrowth kinetics (r_wt_ - r_x_, black) or as survivorship coefficients for competitive aging (*S*, red). Each data point is the average of at least six technical replicates of monocultures or competitions in an independent experimental replicate. Horizontal lines are the mean of each data series. Stacked bars below the plot indicate the number of replicate plates per sample; solid fraction indicates the number of significant samples compared with the WT distribution (*p*<0.01, *t*-test; each experimental batch has 31 reference and at least six deletion samples).

To quantitatively contrast the replicability of both experimental approaches, we fitted decay curves from the outgrowth kinetics experiments to an exponential model. We then compared the difference of adjusted exponential death rates of wild-type and mutant strains from monoculture aging to the *S* parameter obtained from competitive-aging. We observed that both methods performed similarly when comparing the technical variation within each of the plate replicates; there was no significant difference in the typical standard error of the mean in outgrowth kinetics and competitive aging (Supplementary Figure S1). Likewise, when looking at the correlation of quantitative data resulting from independent replicates, we found equivalent correlation coefficients between replicates of each of the two approaches, showing that competitive-aging is as replicable as the outgrowth-kinetics method (Supplementary Figure S1). We note that *gln3*Δ was not included in one of the experiments, and that the *ras2*Δ was atypically noisy in one of the outgrowth-kinetics replicates (not shown); hence these strains were conservatively excluded from this analysis to prevent overestimation of the intra- and inter-batch variability of the outgrowth-kinetics approach. Together, these results indicate that the competition-based method is highly and at least as replicable as the established method, despite inherent variation of different experimental batches.

We looked closer into the distribution of effects of all deletion mutants in five replicates, as determined by the two experimental approaches (**Figure 2C**). Competitive-aging screening showed higher dynamic range but also higher variability. Deletion of *rim15*Δ consistently showed strong short-lived phenotypes with both approaches, while milder short-lived phenotypes of *hap3*Δ, *ste12*Δ, *msn4*Δ, and *msn2*Δ were also scored in most of the plate replicates. Virtually all long-lived phenotypes were correctly scored by both approaches, except for one replicate of *swr1*Δ by the OD-kinetics method. The short- and long-lived phenotypes of *atg1*Δ and *rps16a*Δ, respectively, were not recapitulated by any of the two methods, most likely due to differences in our experimental conditions or strain background.

Having multiple reference samples (WT_RFP_+WT_CFP_) in each 96-well plate improves the fit of the model, but necessarily reduces the number of samples in large-scale analyses, which usually require running many batches in parallel. To maximize experimental throughput without losing quantitative resolution, we used this experiment to estimate the optimal number of reference samples per plate to estimate of *S*. To do so, we quantified how the number of WT-reference samples (from 1 to 31 in this experiment) used to fit *S* = 0 affected the variation of mutant’s *S*. We observed that both the average standard deviation of *S* and the confidence intervals of the fit of *S* showed a steep decline from one to five reference samples, after which variation kept decreasing with diminishing returns (Supplementary Figure S2). Thus, we conclude that including 6 to 10 reference samples is enough to provide a robust description of relative-CLS phenotypes in large-scale genetic analyses of mutants, environments, and their interactions.

Finally, we used this data set to evaluate the influence of the growth rate of the mutants in the context of our competitive-aging method and multiple-regression model, which includes parameter *G*. Importantly, we observed that the model correctly identified mutants with growth defects, as measured by independent culture outgrowths (Supplementary Figure S3); hence, this parameter may enhance a more accurate quantification of relative survivorship of deletion strains. To compare the estimation of the effects on survivorship upon introduction of the *G* parameter, the data was fitted assuming *G* = 0 in all mutant-wild-type competitions; we observed that the difference in the estimation of *S* when *G* was included was modest, but mostly explainable by *G*. As expected, the relative survivorship of slow-growth mutants (*rim15*Δ, *gln3*Δ, and *ras2*Δ) was underestimated when differences in growth rate were not taken into account (Supplementary Figure S3). These results indicate that multiple-regression model optimizes data analysis, especially in screens including mutants with impaired exponential growth.

### Competitive-quantification of CLS under different conditions successfully describes gene-drug interactions

Lifespan is a complex trait determined by different cellular pathways, hundreds of genes, and environmental variables^2,31^. A current challenge in the field is to understand how different factors are integrated with one another to control cell survivorship. Our competitive-aging method provides high-resolution and replicable data, which enables an accurate quantitative description of CLS-phenotype interactions. As a proof of principle experiment using our optimized experimental design and data analysis, we screened for gene-environment interactions in an array of knockout mutants aged with and without the lifespan-extending drug metformin.

We confirmed that the CLS of WT reference samples of yeast increased significantly from a half life of 10.7 to 15.5 days when treated with 40mM metformin (**Figure 4A**; *p*<10^-9^, Wilcoxon rank sum test). Next, we used our competitive-aging approach to measure the CLS of an array of 76 knockout strains aged with or without metformin (random selection of strains from the yeast deletion collection). In both conditions, we observed high quantitative correlation of the phenotypes between replicate plates in the same experiment and between two independent experiments (Supplementary Figure S4). We also confirmed that parameter *G* predicted actual growth rates and resulted in a modest correction of *S*, as expected (Supplementary Figure S5).

**Figure 4.**
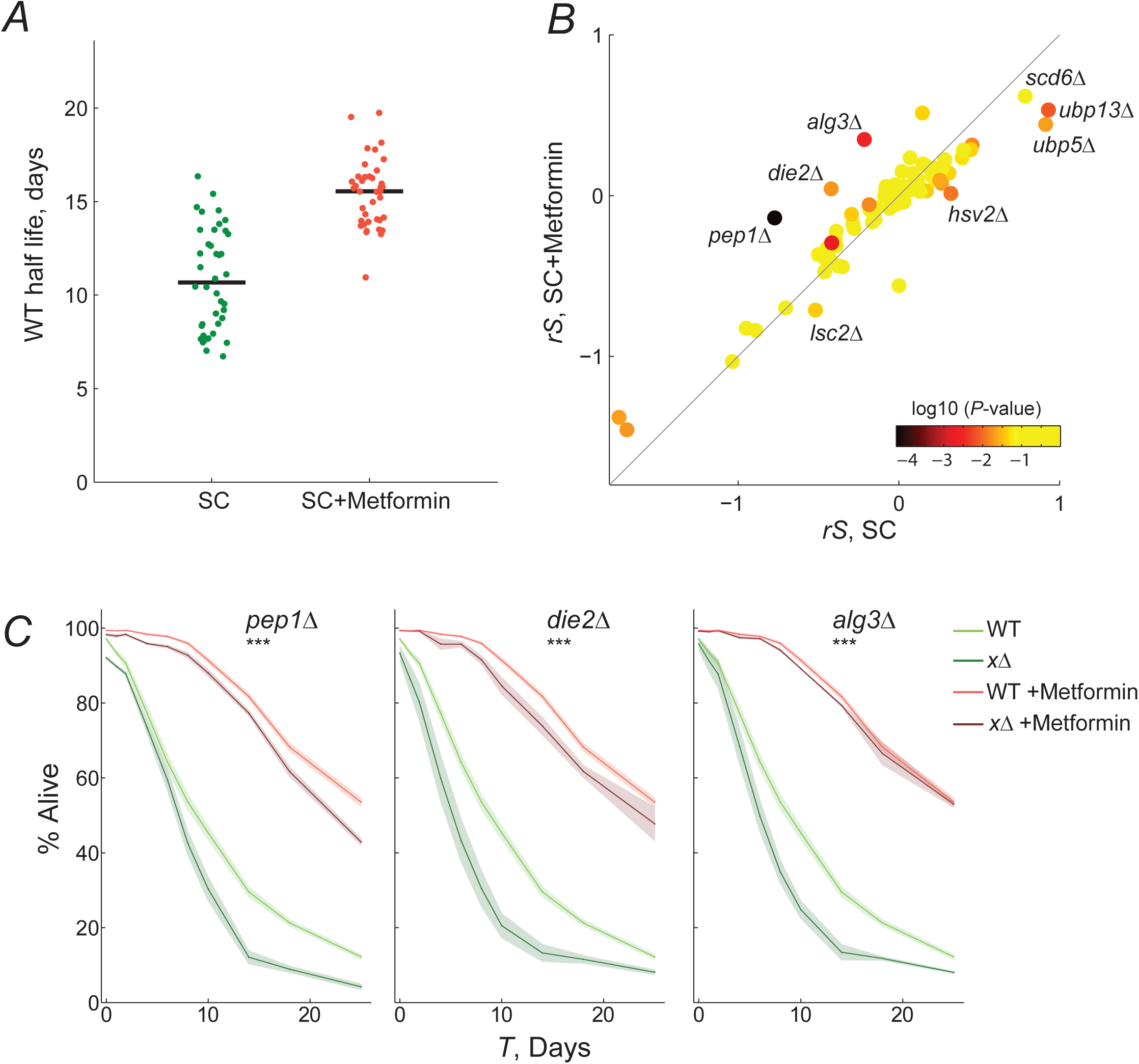
Identification of gene-drug interactions by competitive-aging screening. ***A***, Half life of WT-reference samples with (*n*=39) or without metformin (*n*=40). ***B***, Scatter plot comparing the CLS phenotypes of 76 gene-deletion strains with or without metformin; the average rescaled relative survivorship *rS* shown of four replicates is shown. Color scale indicates the *p-*value of paired t-tests between the *rS* in SC and SC with metformin. ***C***, Gene-drug interactions confirmed by live/dead staining. Survival curves of wild-type (light colors) and gene-deletion (dark colors) strains in nominal SC medium (green) or SC with metformin (orange); 95% CI were calculated from at least three replicates (shaded area). The interaction between metformin and all three gene-deletions was scored significant by two-way ANOVA tests of the death rates (\*\*\**p*< 0.01).

A direct quantitative comparison of CLS phenotypes under both conditions is shown in **Figure 4B** (see Supplementary Figure S6 for data rescaling). Most samples were found to fall close to the diagonal; namely the phenotypic effect of the knockout relative to WT was similar under both conditions. The phenotypes of 21 of the 76 knockouts (27%) were significantly different when treated with metformin (Supplementary Figure S7; *t*-test, *p*<0.05). Mutants with significantly-different phenotypes are potential gene-drug interactions, especially those instances with large deviations from the expectation. In many cases, we observed that metformin alleviated or even reverted the short-lifespan effects of the gene deletion, with *pep1*Δ, *die2*Δ, and *alg3*Δ being the most extreme instances. In only one mutant, *lsc2*Δ, the drug significantly aggravated the relative short-lived gene-deletion phenotype. On the opposite scenario, the nominal long-lived phenotype of certain mutants was rendered neutral or closer to neutral with the metformin treatment (eg. *hsv2*Δ, *ubp13*Δ, and *ubp5*Δ).

We aimed to validate some of the potential gene-drug interactions identified in our screen, focusing on eight mutants with large deviation from the expectation. First, we measured the CLS of wild-type and gene-deletion strains with or without metformin, using the outgrowth OD-kinetics approach (Supplementary Figure S7). A clear effect of metformin treatment on the CLS phenotype was evident for *pep1*Δ, *die2*Δ, and *alg3*Δ; in these cases the short-lifespan phenotype of the mutant was suppressed or alleviated by metformin. We carried out live/dead staining of stationary-phase cells on these three strains, which confirmed gene-drug interactions (**Figure 4C**). The identified interactions, with specific quantitative information on the magnitude and sign of the effects, indicate a functional association between protein glycosylation and metformin treatment, providing novel information to understand the mechanisms of longevity by metformin in yeast. Together, results in this section show that competitive-aging yields accurate and replicable large-scale CLS data in a straight-forward manner to shed light on the mechanisms of pharmacological interventions that extend lifespan.

## DISCUSSION

Genetic analysis of the CLS of budding yeast has led to the genome-wide identification of genes involved in aging; recent efforts have sought to describe interactions between genetic and environmental modulators of the phenotype (Garay et al., 2014; Smith et al., 2016; Campos et al., 2018; Jung et al., 2018). A previous report from our group showed advantages of using a competitive-aging approach, in which fluorescently-labeled strains in co-culture provide high resolution in parallel setups (Garay et al., 2014). Competitive-aging has also allowed scoring gene-environment interactions at the genomewide level, specifically interactions with dietary restriction (Campos et al., 2018).

Here, we have presented an optimized model to calculate the relative survivorship of deletion strains, taking into account the possible confounding effects coming from growth-rate differences and systematic batch effects. It is certain that major findings in our previous reports would hold given the relatively mild contribution of the *G* and *C* parameters in most samples (the experimental design of such studies precludes fitting data to the model herein presented). Nonetheless, the optimized procedure herein presented is a more exhaustive description of the actual competitive-aging setup, which could improve the scoring CLS phenotypes in specific cases, particularly strains with strong growth defects or experiments with strong batch effects. In addition, we directly compared the performance of a competitive-aging setup to an established approach (Murakami et al., 2008; Jung et al., 2015). Finally, using this enhanced method and data analysis in a proof-of-principle experiment, we were able to unravel significant gene-drug interactions in an array of gene-deletions strains subjected to the lifespan-extending drug metformin.

Early CLS genome-wide screens were based on large pools of gene deletions followed by molecular-barcode hybridization or sequencing (Fabrizio et al., 2010; Matecic et al., 2010; Gresham et al., 2011). These studies provided important insight into which genetic factors mediate stationary phase survival, such as autophagy, vacuolar protein sorting, and regulation of translation. However, the high rates of false positives— specially in the cases of long-lived phenotypes—and low overlap among the sets of genes from different studies (Smith et al., 2016) suggest that systematic errors in barcode detection or major experimental batch effects result in poor experimental replicability. On the other side of the spectrum, an ingenious outgrowth-kinetics approach of yeast monocultures increases the feasibility of percent-survivorship estimates, compared to the conventional colony-forming units method (Murakami et al., 2008); but throughput is still limited with this approach. To overcome this limitation, Jung and co-workers scaled-up this strategy using monocultures in multi-well plates, whereby more strains can be tested in parallel (Jung et al., 2015, 2018). There is still the issue that mild environmental variation can affect separate cultures differently, that could lead to low reproducibility (Burtner et al., 2011); for instance, strains may reach stationary phase at different times after inoculation. In this regards, competitive-aging provides a direct phenotypic comparison and, arguably, more consistent results, given that the mutant population of interest is aged with an internal reference strain under the exact same microenvironment. Importantly, competitive-aging can also be carried out in multi-well plates, enabling high-throughput experimental setups.

One of the inherent drawbacks of the competitive-aging method is that no absolute death-rate information is provided, which can be easily solved thorough characterization of the wild-type strain in monoculture, as we have shown. Likewise, the dynamic range of this method depends on the actual death rate of the reference strain used. For instance, quantitative phenotypic descriptions are limited if *S*>>0; in other words only semi-quantitative estimates are possible for extremely long-lived strains (*S* ≈ *r_wt_*). The use of different reference strains, eg. specific gene deletions with known long-lifespan phenotypes, may help to overcome this limitation. In addition, just as most population-based CLS methods, competitive aging depends on cells being able to re-enter the cell cycle, which can only be distinguished from actual death using outgrowth-independent methods, such as live/dead staining.

We have illustrated the potential of competitive-aging screening by characterizing an array of yeast deletion strains exposed to metformin. A number of lifespan-extending pharmacological interventions are already being tested for age-related diseases in humans, even when the mechanisms underlying their beneficial effects are frequently not fully understood (Mullard, 2018; Zimmermann et al., 2018). Given that lifespan is a complex phenotype, identifying conserved gene-drug interactions could shed light on the modes of action and to pinpoint genetic modifiers of the drug’s effects (Zimmermann et al., 2018). By screening an array of 76 gene deletions aged with metformin, we found a number of cases in which metformin buffers both short- or long-lifespan mutant phenotypes. The interacting genes suggest a role of protein glycosylation and protein homeostasis, which is in line with previous evidence showing that metformin alters glycation, protein transport, and protein degradation in yeast (Borklu-Yucel et al., 2015; Kazi et al., 2017). Our results also uncovered interactions between metformin and genes involved in mitochondrial function (*erp6*Δ, *lsc2*Δ*, ema35*Δ, and *ylh47*Δ), which is a known player in the cellular response to metformin (Borklu-Yucel et al., 2015; Stynen et al., 2018). It remains to be addressed how these proteins are specifically related to the known response involving the mitochondrial electron transport chain and homeostasis of copper and iron.

Competitive-aging can readily be adapted to screen double mutants at large scale and to score genetic interactions underlying CLS phenotypes. Genetic interactions (epistasis) are a powerful tool to describe the architecture of phenotypes and the functional relationships of different genetic pathways (Segrè et al., 2005; Collins et al., 2007; Onge et al., 2007; Phillips, 2008; Kuzmin et al., 2018). While epistasis-network analyses in yeast have granted deep knowledge of the genetic landscape of mitotically-active, proliferating cells, less is known about how genetic interactions shape the genetic architecture of post-mitotic survivorship. Large-scale genetic analysis of double-mutants aged in competition with their single-mutant references is an attractive experimental setup to identify interactions among different genes and pathways underlying CLS in yeast. Our competitive-aging screening method and quantitative analysis provide a powerful systematic tool to shed light on the complex genetic, environmental, and pharmacological wiring of aging cells.

## Acknowledgements

We are grateful to Erika Cruz-Bonilla, Nelly Selem, and Judith Ulloa for critical reading of the manuscript. This work was funded by the Consejo Nacional de Ciencia y Tecnología de México (CONACYT grants CB-2015/164889 and PN-2016/2370). J.A.A.-R. received a doctoral fellowship from CONACYT (#264529). The funders had no role in study design, data collection and analysis, decision to publish, or preparation of the manuscript. This manuscript has been released as a pre-print at BioRxiv (Avelar-Rivas et al., 2019).

## Author Contributions

Conceived and designed the study: JAA-R, AD. Performed the experiments: JAA-R, MM-F, AJ-R, SEC. Developed and implemented the model: JAA-R, EG, NS. Analyzed the data: JAA-R, SEC, AD. Wrote the paper: JAA-R, AD. Acquired funding: AD. All authors read and approved the final version of the submitted manuscript.

## SUPPLEMENTARY ITEMS (ZIP)

**Note S1.** Development and implementation of a linear model to calculate relative survival of mutant strains in competitive-aging cultures (PDF)

**Data S1.** Array of mutants used for gene-drug interactions, with their relative survivorship, rescaled survivorship, and relative growth rates (XLSX)

## Supplementary Note 1. Development and implementation of a linear model to calculate relative survival of mutant strains in competitive-aging cultures

To obtain relative survivorship of competitive-aging cultures we modeled WT and mutant populations with initial sizes *N_wtT0_* and *N_xT0_* and declining trough time in stationary phase at constant exponential rates *r_wt_* and *r_x_*. After *T_i_* days, the sizes of the remaining viable populations are 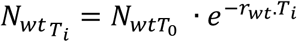 and 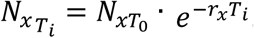. In every outgrowth sampling, we inoculated small volumes of this aged co-culture onto fresh medium, in which both populations resume cell division and grow exponentially at rates *g_wt_* and *g_x_* At any outgrowth, the population sizes after *t_j_* hours are 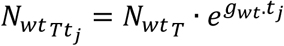 and 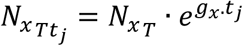. Given that both strains are aged and grown in co-culture, the relationship between the sizes of their populations at any time, *t_j_*, during the outgrowth *T_j_* is:

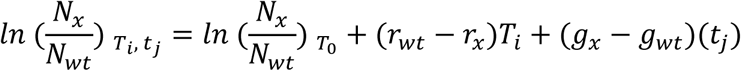

As shown in Figure S1, the measurements of *CFP* and *RFP* signals are proportional to *N_wtTt_* and *N_xTt_*, therefore, the experimental measurements of each well, *w* are equal to the linear equation:

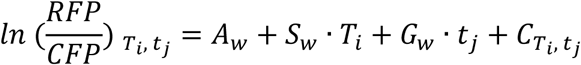

We run a multiple linear regression where we fit the data obtained from measuring bulk RFP signal (*RFP*), and bulk CFP signal (*CFP*) in the outgrowth cultures in fresh medium sampled by inoculating from aging stationary phase co-cultures. The first step is to subtract auto-fluorescence *RFP*_raw_ from WT_CFP_ as a function of the *OD_600_* from all *RFP*_raw_ measurements; the same is done for *CFP*_raw_ from WT_RFP_. We keep outgrowth measurements for days in which we observe an increase in both RFP_raw_ and *CFP*_raw_. This leaves us with the *RFP* and *CFP* quantities that should be a function of the death and growth rates (*r* and *g*). For each measured well, *w*, we expressed such relationship using the model 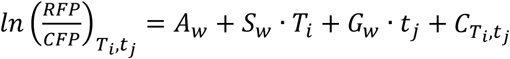 in which *A_w_* reflects the proportion 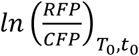 at the beginning of the experiment, this number will depend on the amount of Δ*x*_RFP_ and WT_CFP_ strains that are mixed together when the aging co-cultures are set up. The relative survivorship is modeled in *S_w_* = (*r_wt_* - *r_x_*), which is expressed in terms of the number of the time *T* (in days) that the co-culture has been in stationary phase. Meanwhile the difference in growth rates of both strains is modeled in *G_w_* = (*g_x_* - *g_wt_*), which is expressed in terms of the time *t* (in hours) within every outgrowth. Finally, the model takes into account the batch error of each plate read, as *C_Ti,tj_*.

To fit the data, we generate a system of linear equations. For well *w*_1_, there are as many equations as the total number of measurements of that well across the entire experiment. For instance, day number one,*T*_1_, has the following equations:

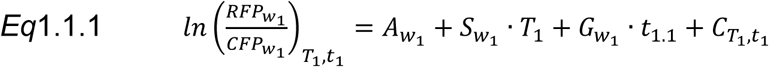

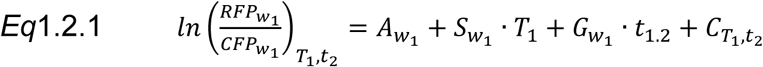

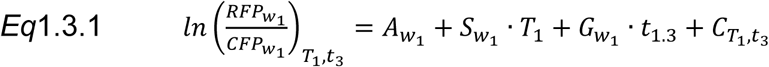

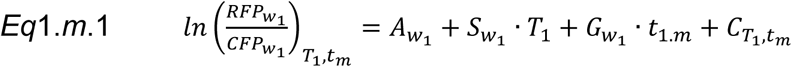

In these equations, the term 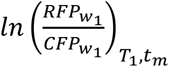 is the empirical determination of the logarithmic quotient of both fluorescent signals at each measurement of the outgrowth in exponential phase. *A_w_*_1_ is the same variable for all equations of *w*_1_, likewise for S_w1_ · *T*_1_ because the relative death rate of the well (*S_w_*_1_) is the same and is multiplied by the same number of days of the experiment, *T*_1_. By definition, the first day of measurement is considered time *T*_1_ = 0 for all plates and wells. One of the two terms that change between these equations is, *G_w_*_1_ · *t*_1.*m*_. Although the relative growth rate is the same for each well throughout the entire experiment, the time at which the measurement is done changes and that is reflected in the different coefficients that these equations will get for those terms. For example, if measurements are done every 1.5 hours and the exponential phase starts at the third measurement, coefficients would be *t*_1.1_ = 3; *t*_1.2_ = 4.5; *t*_1.3_ = 6. Finally, the other term that changes in these equations is *C_T_*_1,*tm*_ which reflects that there are batch effects influencing measurements; such effects appear any time a plate is read, and so it has similar effects in all of the wells within the same plate. For that reason *C_T_*_1,*tm*_ is different at each measurement of a well, but is shared in all wells from the same plate read. Therefore, the measurements of different wells in one plate with *k* wells lead to equations:

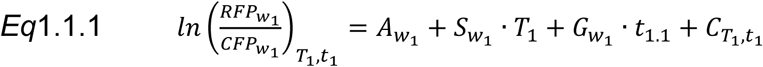

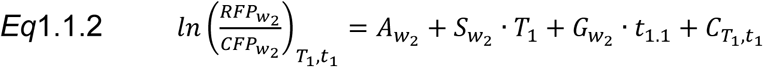

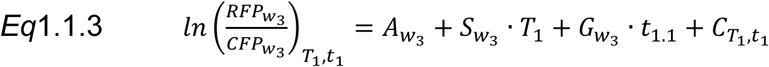

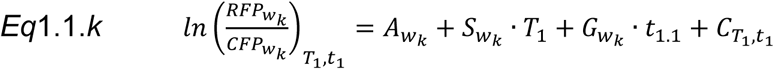

The same wells measured the second day will have the equations:

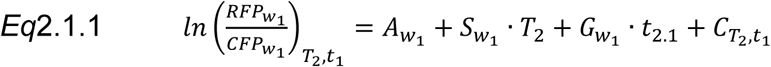

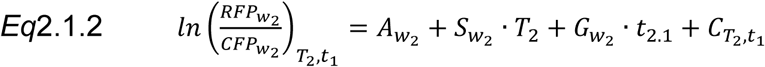

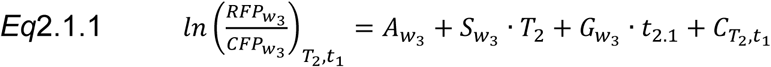

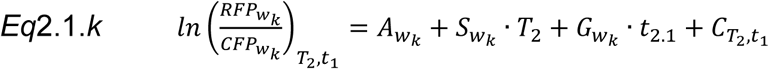

If the second outgrowth occurs after one day then *T*_2_ = 1. It is worth pointing out that this *t*_2.1_ is different from *t*_1.1_ because the outgrowths of each day are not measured at identical points in time. For example, it could be that *t*_2.1_ = 3.6 and *t*_2.2_ = 4.1.

We verified the quality of the data and set some restrictions. For instance, in every valid measurement both fluorescent signals must be above the maximum auto-fluorescence detection, as indicated by wells with monocultures of WT_CFP_ or WT_RFP_ strain with auto-fluorescence of the opposite signal. This step removes all monoculture wells from the system of equations.

Our model fitting has the assumption that WT_CFP_+WT_RFP_ competitions have *S_x_* = 0 and *G_x_* = 0, so deviations from such values in wells *ref*, containing such mixture of strains depend on the well specific variable *A_wref.i_* or to the plate variables *C_Tn,tm_*. To accomplish that, we eliminate *S* and *G* from the reference-well equations, so every measurement from those wells will feed the system of equations as follows:

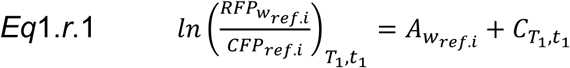

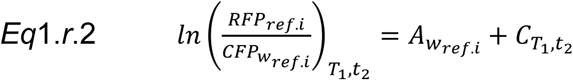

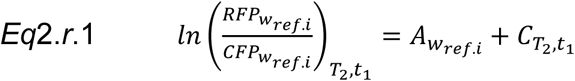

To find an approximation to the solution of the entire set of equations, we built a predictor matrix that contains all the valid measurements of the plate and the logarithmic ratio of the actual experimental measurements as the responses vector. In this example, we show the matrix for wells *w*_1_,*w*_2_, *w_ref_*, and *w_k_* outgrown at days *T*_1_, *T*_2_, and *T_n_* with 3 measurements per outgrowth *t_n_*_·1_, *t_n_*_·2_, and *t_n_*_·m_: If there are *k* wells in a plate, the plate was measured *m* * *n* times then there are *k* * *m* * *n* rows in the matrix and there are 3*_k_* + *m* * *n* – 2*N_wref_* variables (or columns) to fit (*Nw_ref_* is the number of wells that contain reference co-cultures); hence, for the previous matrix there are 36 rows and 19 predictor variables.

**Table S1.**
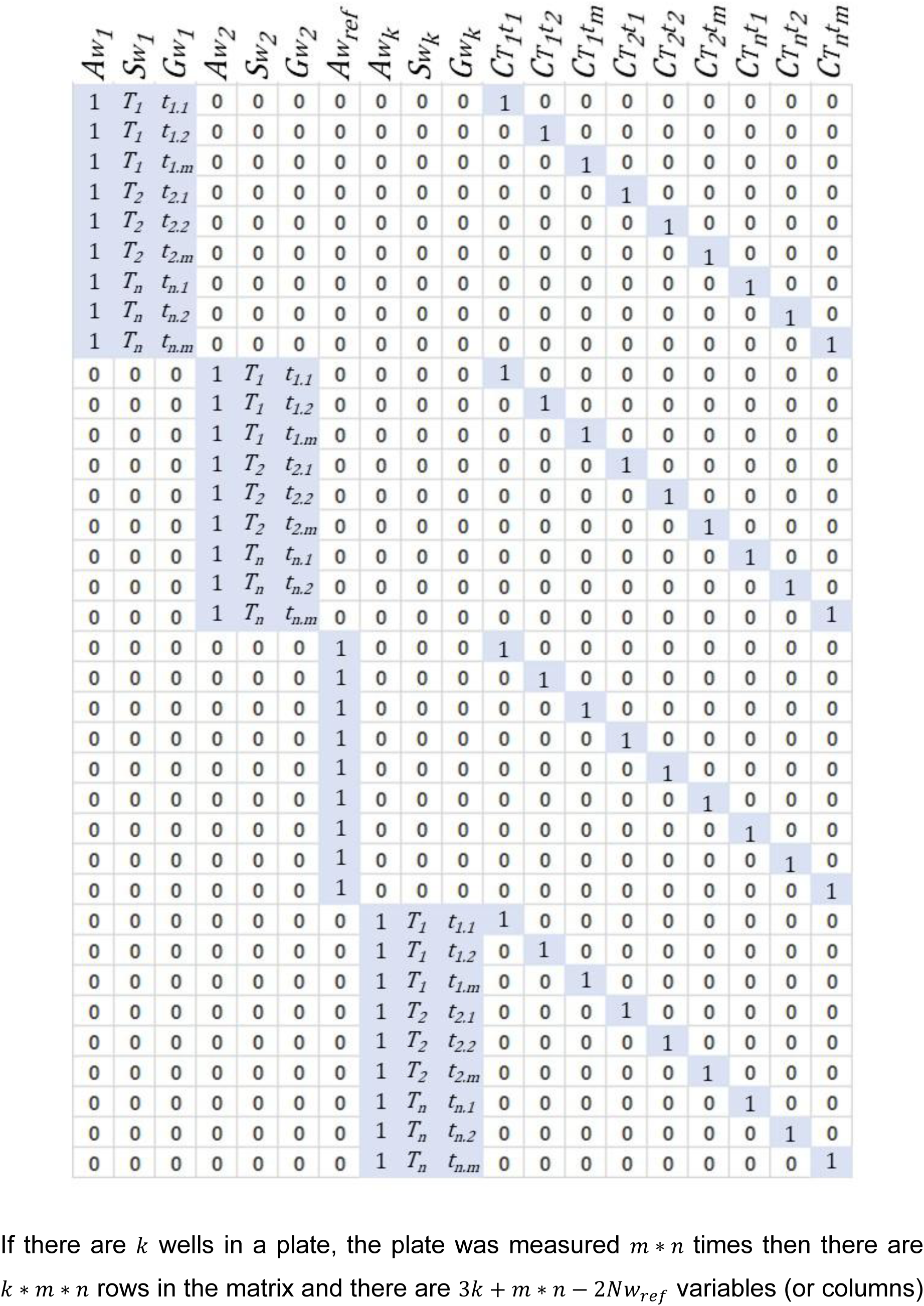
Recipes for media used in this study (PDF)

We applied a multiple linear regression to approximate the solution. The Matlab function *regress* was used, as it provides useful statistics about the fit such as the 95% confidence interval for the coefficient estimates.

Scripts and documentation to run this model are available in GitHub: https://abrahamavelar.github.io/LinearModelCLS/

---

**Table S1.**
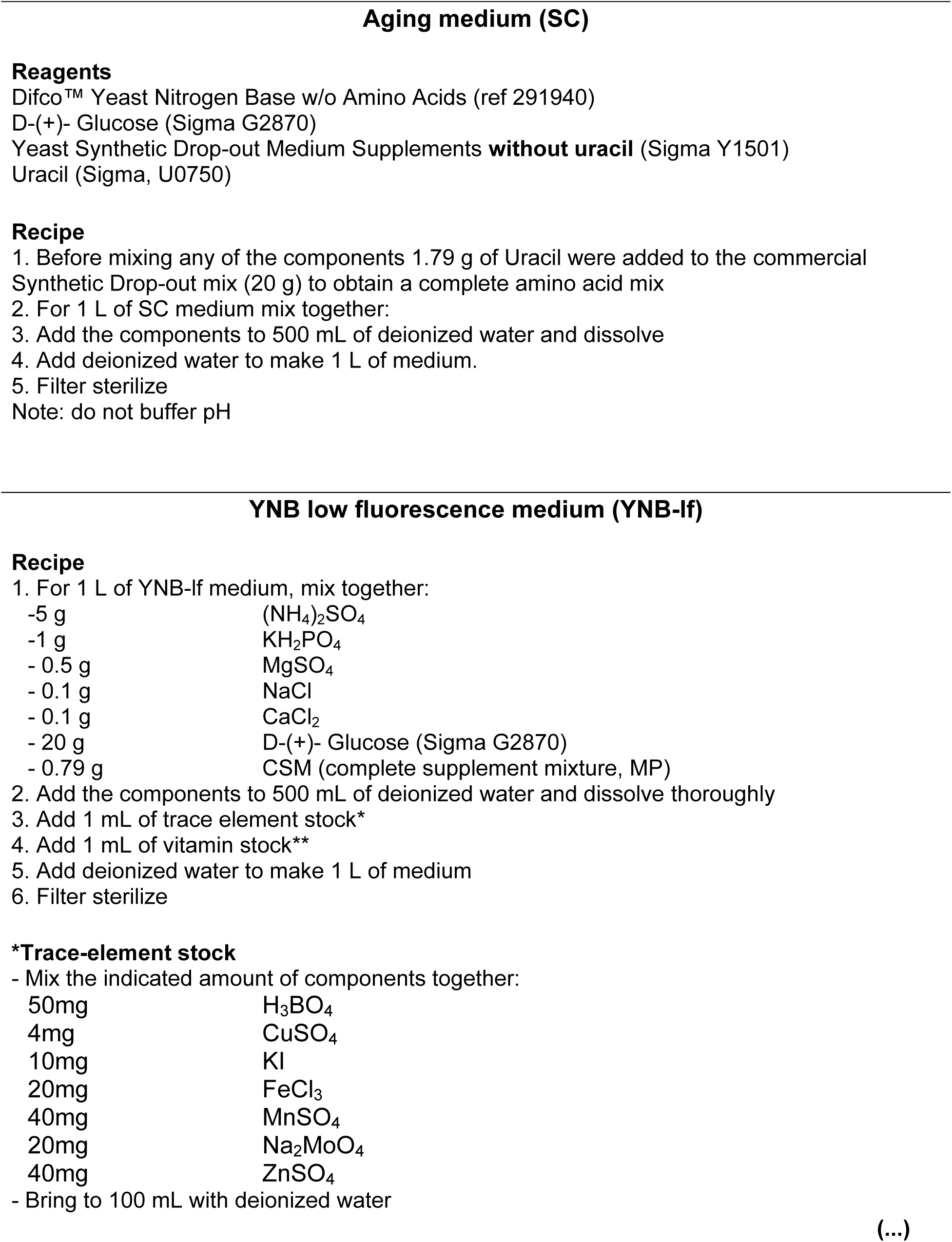
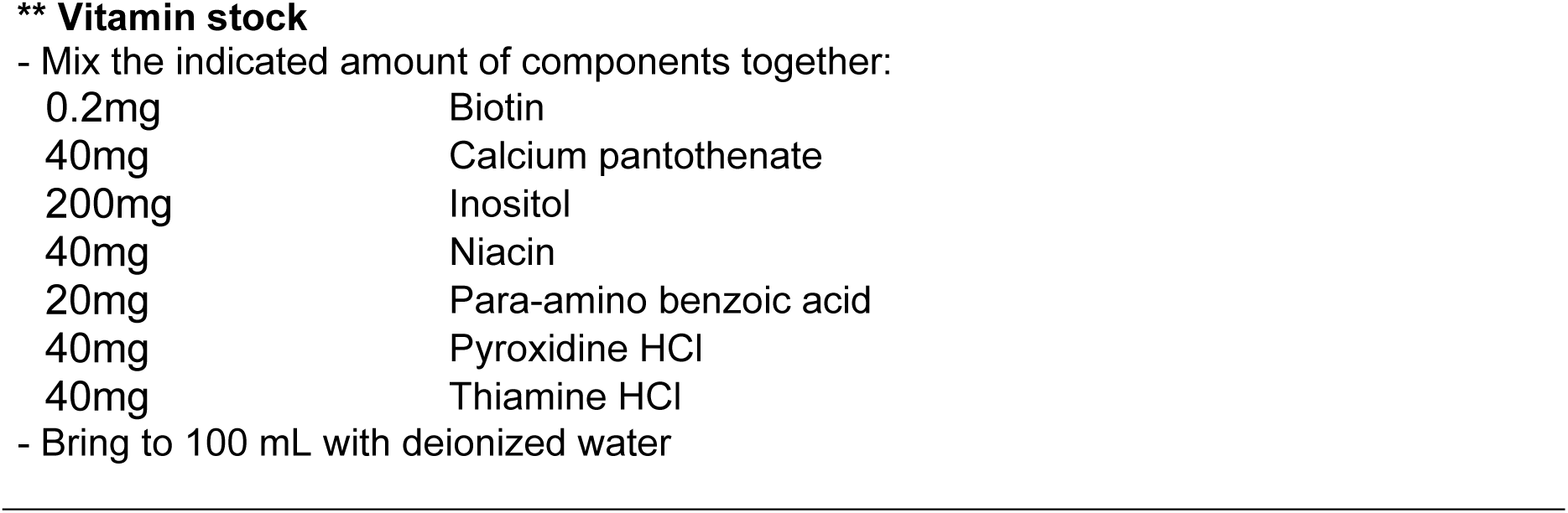
Media recipes

| Aging medium (SC) |  |
| --- | --- |
| **Reagents** |  |
| Difco™ Yeast Nitrogen Base w/o Amino Acids (ref 291940) |  |
| D-(+)- Glucose (Sigma G2870) |  |
| Yeast Synthetic Drop-out Medium Supplements **without uracil** (Sigma Y1501) |  |
| Uracil (Sigma, U0750) |  |
| **Recipe** |  |
| 1. Before mixing any of the components 1.79 g of Uracil were added to the commercial Synthetic Drop-out mix (20 g) to obtain a complete amino acid mix |  |
| 2. For 1 L of SC medium mix together: |  |
| 3. Add the components to 500 mL of deionized water and dissolve |  |
| 4. Add deionized water to make 1 L of medium. |  |
| 5. Filter sterilize |  |
| Note: do not buffer pH |  |
| YNB low fluorescence medium (YNB-lf) |  |
| **Recipe** |  |
| 1. For 1 L of YNB-lf medium, mix together: |  |
| -5 g | (NH<sub>4</sub>)<sub>2</sub>SO<sub>4</sub> |
| -1 g | KH<sub>2</sub>PO<sub>4</sub> |
| - 0.5 g | MgSO<sub>4</sub> |
| - 0.1 g | NaCl |
| - 0.1 g | CaCl<sub>2</sub> |
| - 20 g | D-(+)- Glucose (Sigma G2870) |
| - 0.79 g | CSM (complete supplement mixture, MP) |
| 2. Add the components to 500 mL of deionized water and dissolve thoroughly |  |
| 3. Add 1 mL of trace element stock* |  |
| 4. Add 1 mL of vitamin stock** |  |
| 5. Add deionized water to make 1 L of medium |  |
| 6. Filter sterilize |  |
| **\*Trace-element stock** |  |
| - Mix the indicated amount of components together: |  |
| 50mg | H<sub>3</sub>BO<sub>4</sub> |
| 4mg | CuSO<sub>4</sub> |
| 10mg | KI |
| 20mg | FeCl<sub>3</sub> |
| 40mg | MnSO<sub>4</sub> |
| 20mg | Na<sub>2</sub>MoO<sub>4</sub> |
| 40mg | ZnSO<sub>4</sub> |
| - Bring to 100 mL with deionized water |  |
| 0.2mg | Biotin |
| 40mg | Calcium pantothenate |
| 200mg | Inositol |
| 40mg | Niacin |
| 20mg | Para-amino benzoic acid |
| 40mg | Pyroxidine HCl |
| 40mg | Thiamine HCl |

- Bring to 100 mL with deionized water -

**Supplementary Figure S1.**
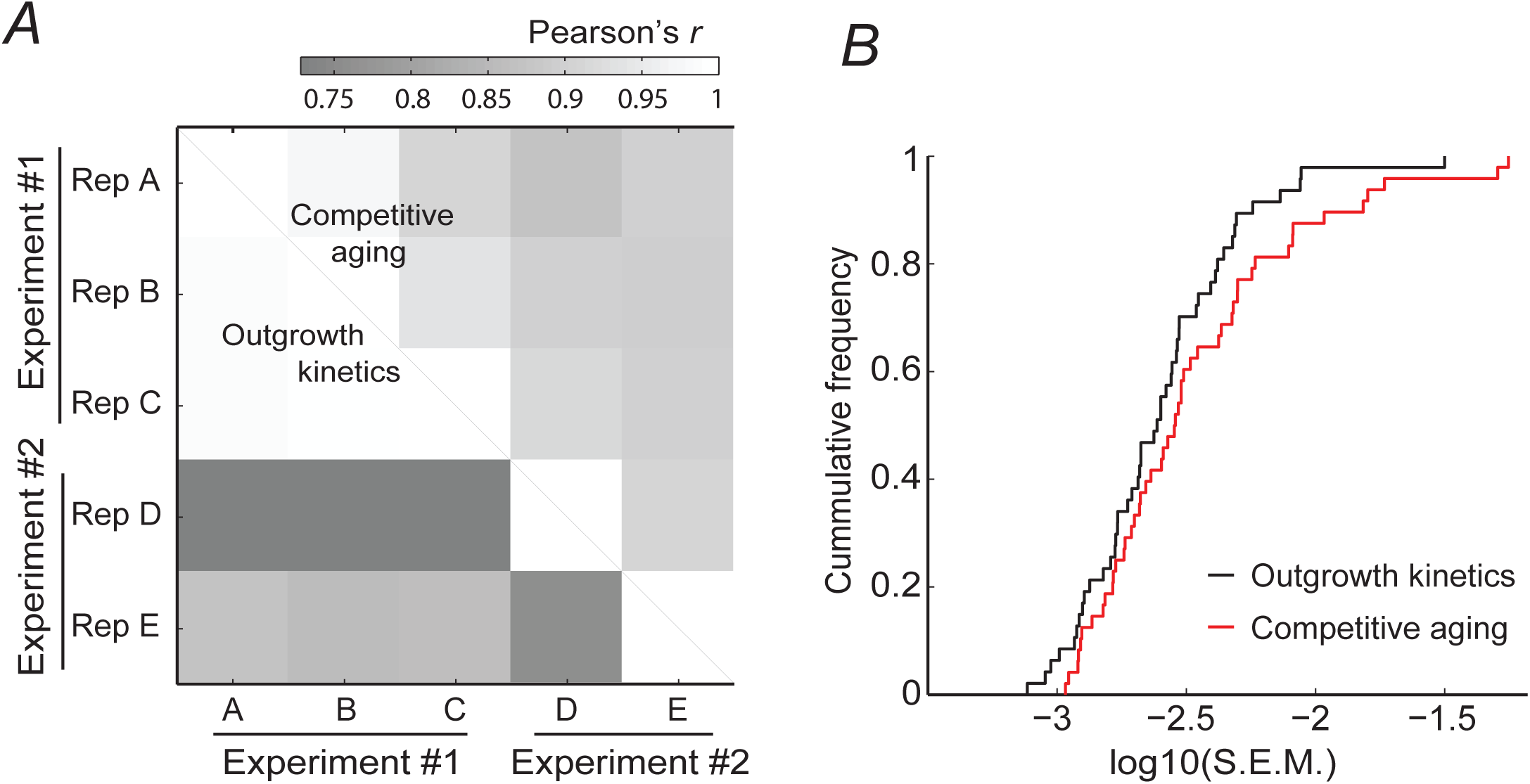
Comparing replicability of CLS measurements from two methods. ***A***, Paired Pearson’s *r* correlation coefficient among the relative survivorship, *S*, from three (experiment #1, plates A, B, and C) or two replicates (experiment #2, plates C and D). Competitive-aging correlations are shown above, while OD-kinetics in monoculture are shown below the diagonal. ***B*,** Cumulative S.E.M. (intra-experiment) of all mutant’s *S*. Data of the five experimental batches are shown in each series (*n*=47 and *n*=48 for monoculture outgrowth kinetics and competitive aging, respectively). The median S.E.M. of *S* from both methods are statistically indistinguishable (*p*=0.154, Wilcoxon rank sum test).

**Supplementary Figure S2.**
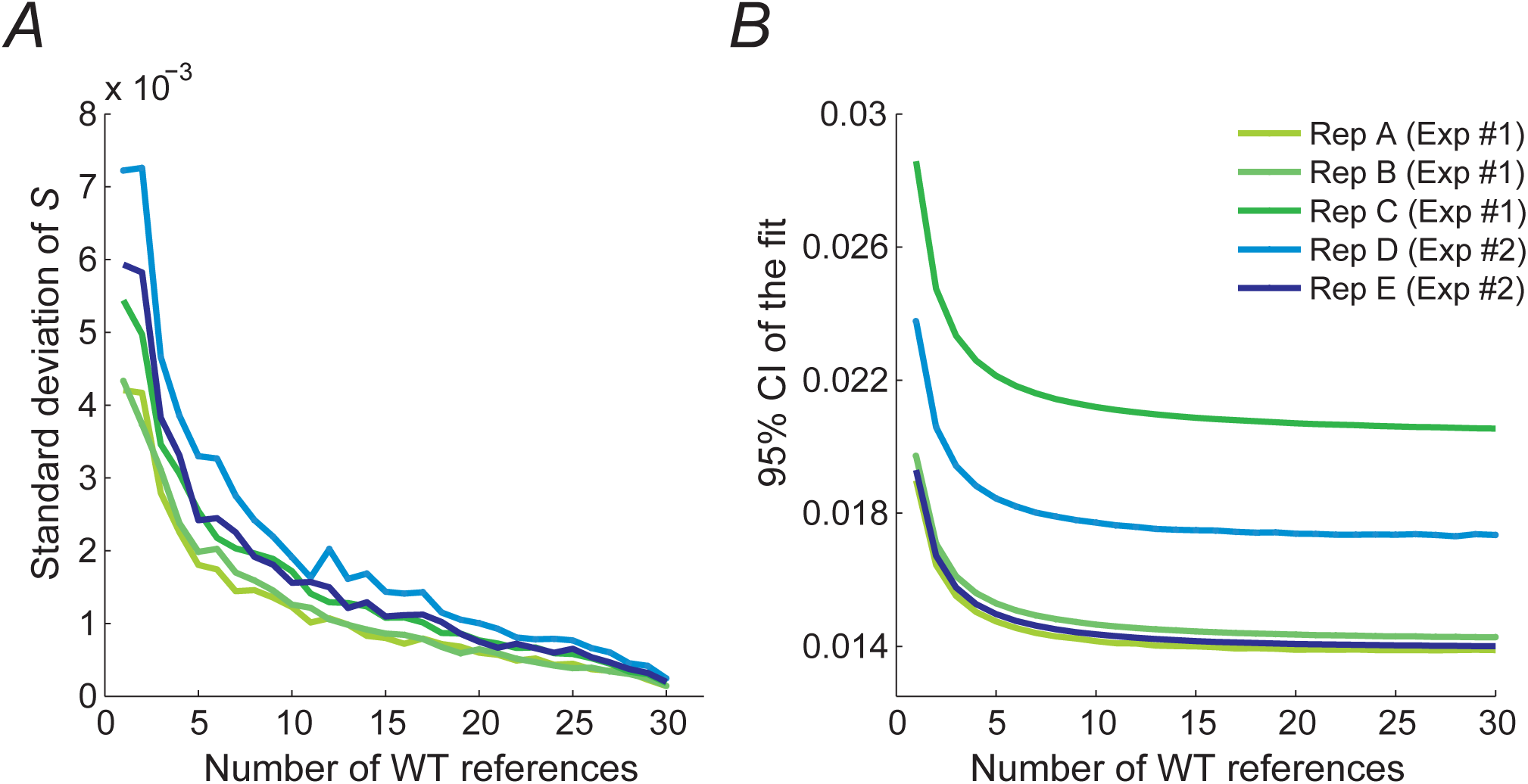
Variation in *S* decreases with the number of reference samples used. Thirty-one WT_RFP_+WT_CFP_ reference competitions were used in each plate. To estimate the effects of increasing number of reference samples, the model was fed with different numbers of reference wells and the reminder control samples were treated as additional mutant samples. For each given number of reference samples, 100 permutations were randomly selected and the selection coefficient (*S*) of all mutant strains was calculated accordingly. ***A***, The variation using different permutations was quantified as the standard deviation of each mutant sample; the average standard deviation of *S* of all mutant samples is shown as a function of the number of reference samples in the model. ***B***, The fit to the model was calculated along with the 95% CIs; the difference between the upper and lower CI bounds is plotted against the number of reference samples added to the model.

**Supplementary Figure S3.**
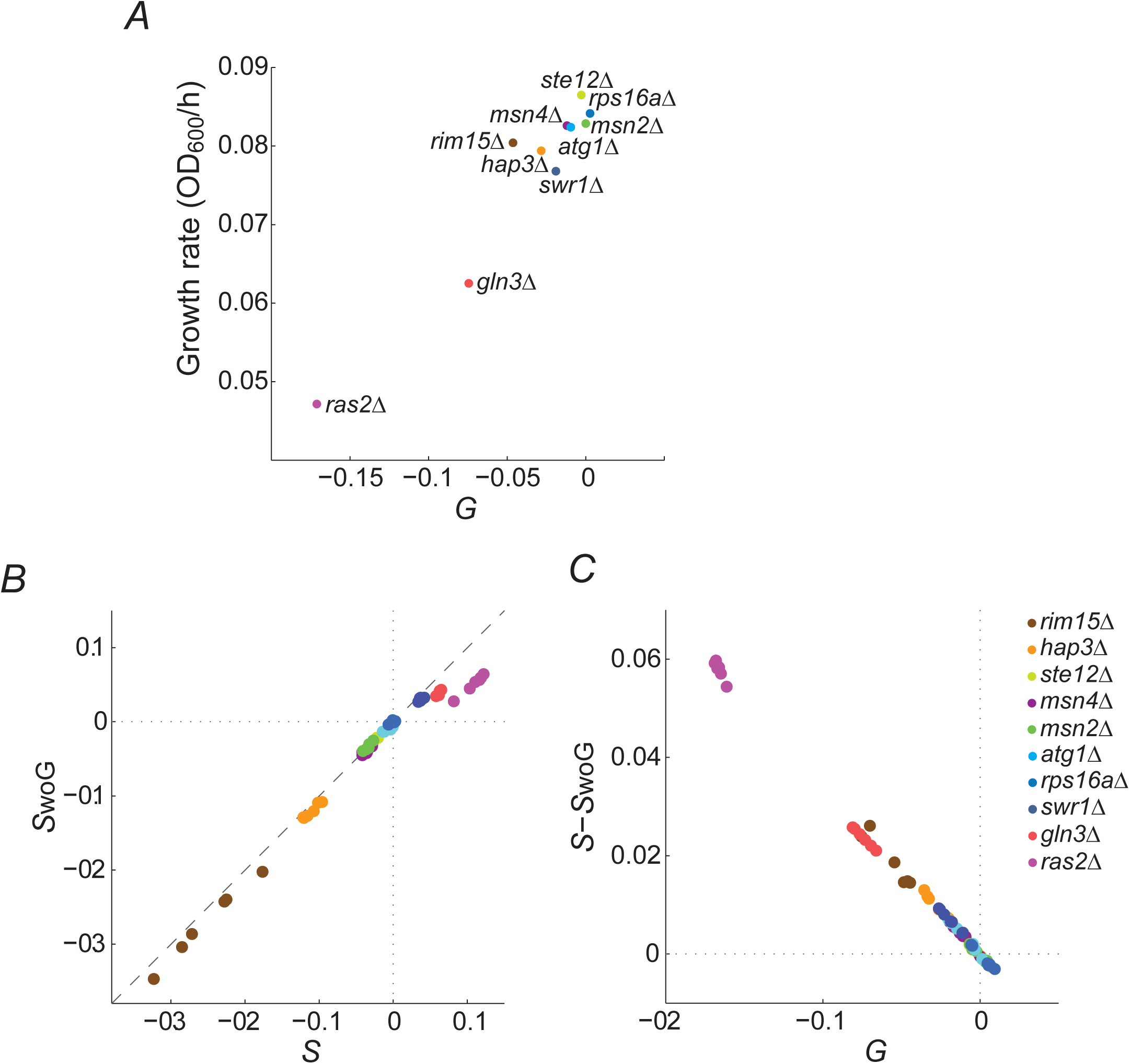
The parameter *G* corrects effects of differential growth rates that otherwise affect relative survivorship, *S*. ***A***, The actual growth rate at exponential phase of gene-deletion strains measured as average increase in *OD_600_* with time (*n=*6-7), compared tothe modeled parameter *G*. ***B***, Scatter plot showing the average *S* value from 3-5 replicates with parameter *G* is included in the multiple linear regression (horizontal axis) and when the regression is run without *G* (*S*_woG_, vertical axis). ***C***, The mild difference between calculations of *S* showed in panel B is mostly explained by changes in the parameter *G*.

**Supplementary Figure S4.**
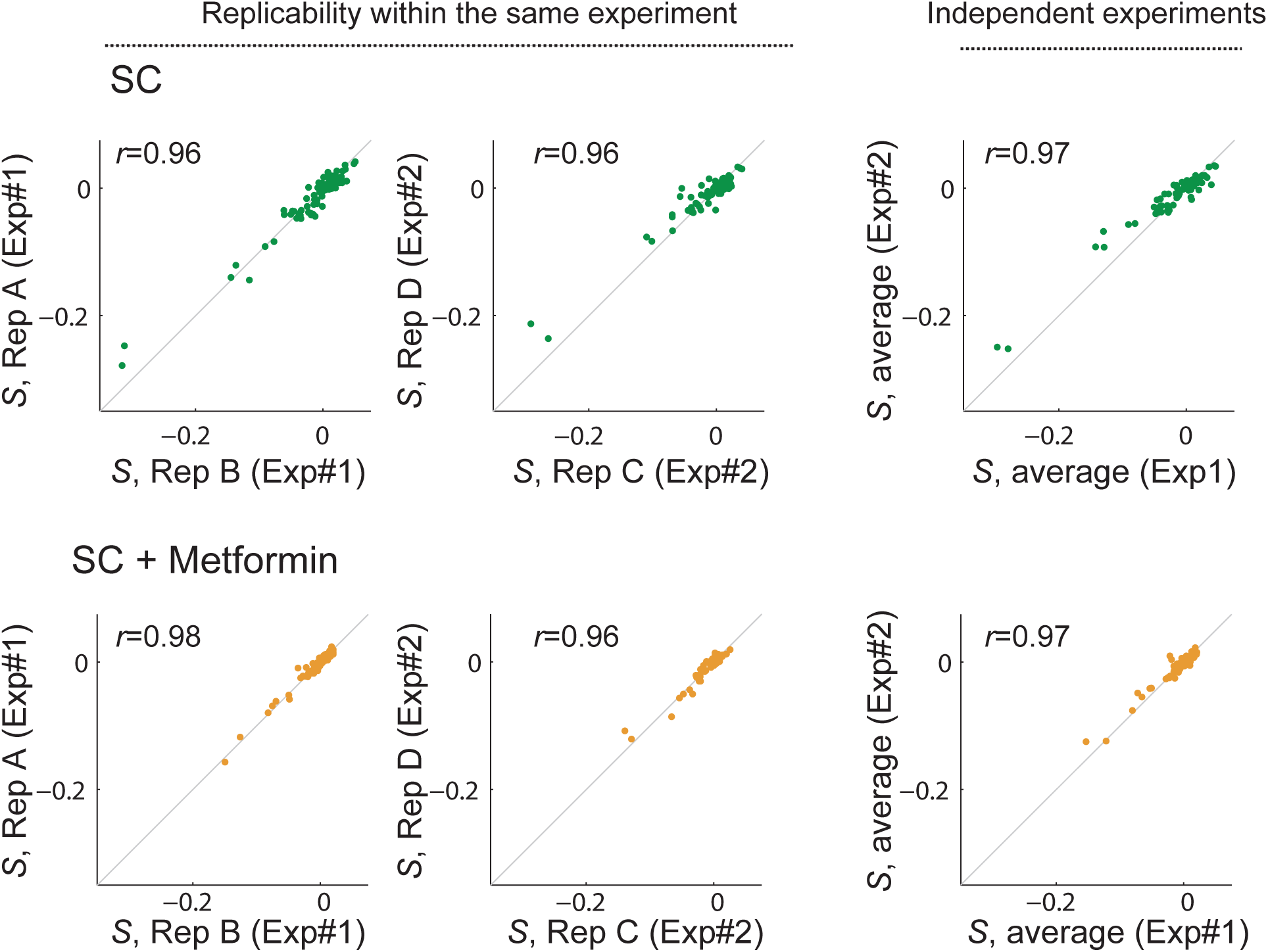
The relative survivorship, *S*, is well replicated within the same experiment (replicates ‘A’ and ‘B’, or ‘C’ and ‘D’) and between different experiments (#1 and #2), both without (green) and with metformin (orange). Left and central panels show single data points comparing two replicate plates, while the right panel shows the comparison of the averaged data of two replicates within each independent experiment. The Pearson correlation coefficients are shown in each panel.

**Supplementary Figure S5.**
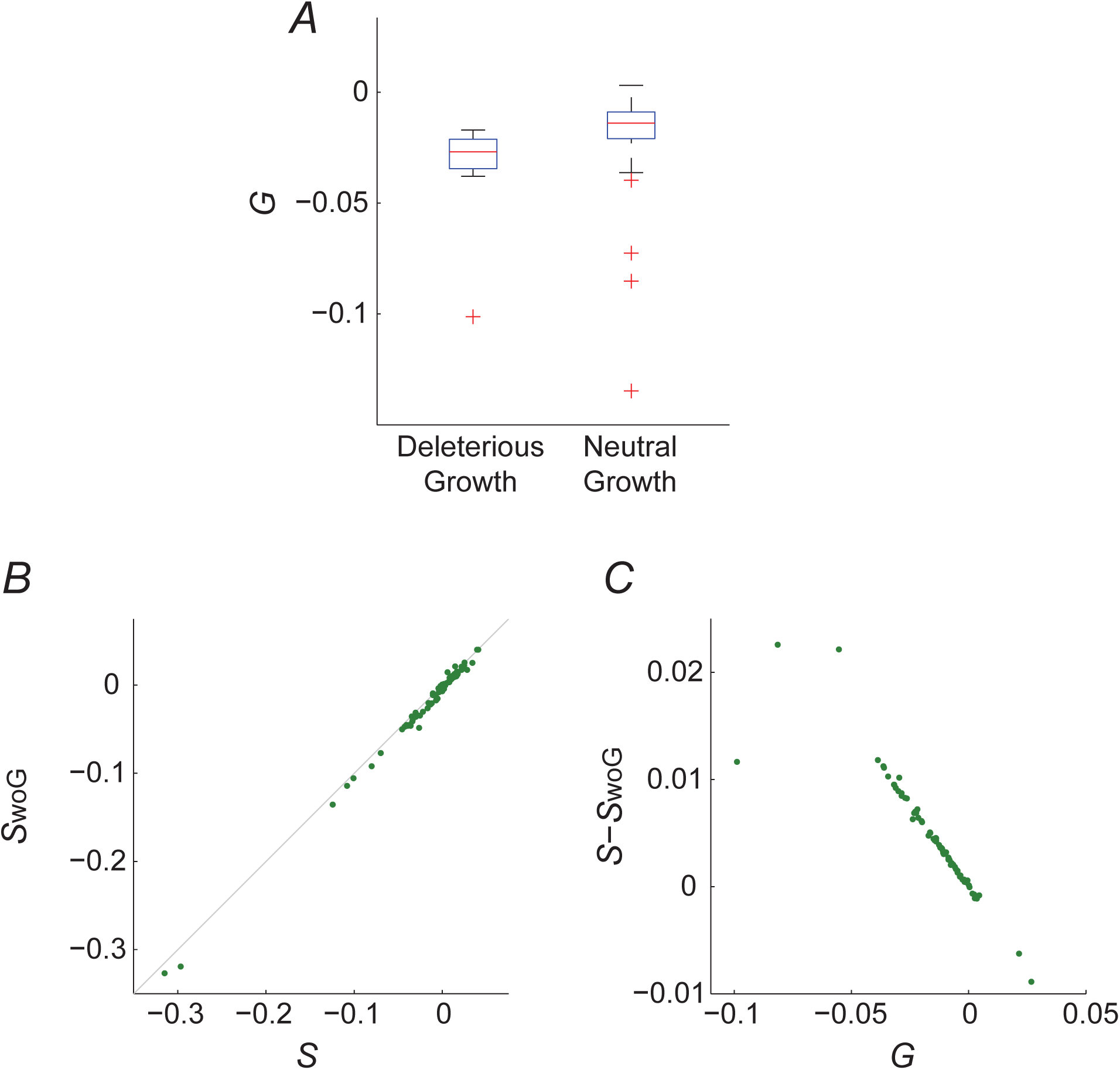
The parameter *G* corrects effects of differential growth rates that otherwise affect relative survivorship, *S*. ***A***, Box plots of the *G* calculated for mutants with reduced fitness or without effects as previously reported (Costanzo et al. 2010). Deleterious knockouts, (fitness *f*<0.95, *n*=10) usually have a more negative *G* value than neutral knockouts without known growth defects (*f*≥0.95, *n*=67) (*p*=0.0014, Wilcoxon rank sum test). ***B***, Scatter plot showing the obtained *S* value with the parameter *G* is included in the multiple linear regression (horizontal axis) and when the regression runs without *G* (*S*_woG_, vertical axis). ***C*,** The difference between calculations of *S* showed in panel A is mostly explained by average changes in the parameter *G*.

**Supplementary Figure S6.**
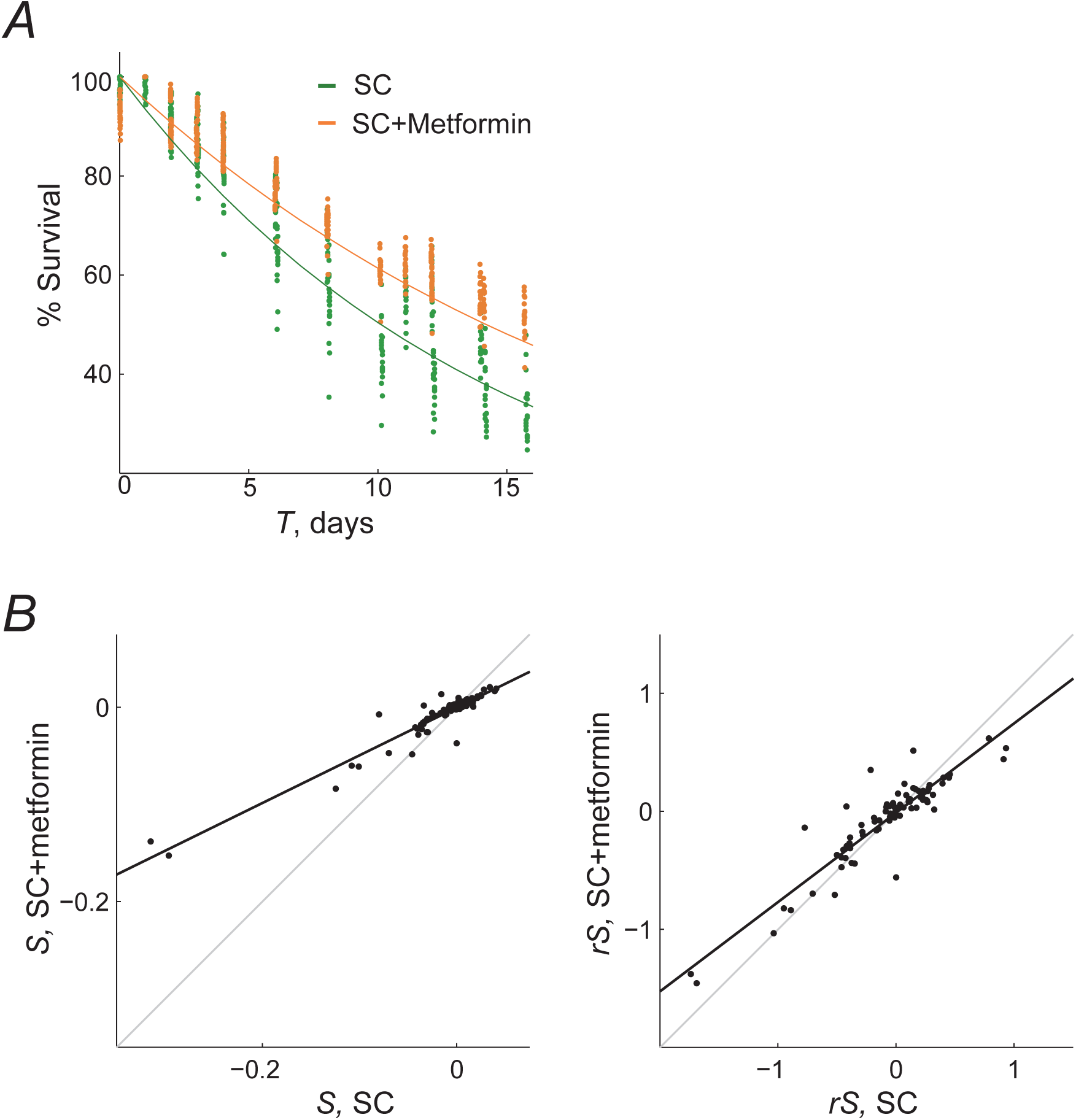
Relative survivorship (r*S*) data rescaling. To quantitatively compare the relative phenotypes between two conditions of different WT phenotype, data were rescaled to a dimensionless relative survival parameter, *rS*. ***A***, The death rates of the WT without (green) and with metformin (orange) were calculated by fitting to an exponential function the percent survival at different days of aging. Data points were from ten replicate samples in four replicate experiments (*n*=39 with metformin and *n*=40 for SC); lines are the average fits. ***B***, The relative survival *S* shows a bias between the two conditions (left); the rescaled survivorship parameter *rS* provides a better comparison under two conditions in which the WT dies at different rates (right). Specifically, relative survivorship was rescaled with the formula 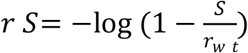 using the average wild-type death rate obtained in each plate.

**Supplementary Figure S7.**
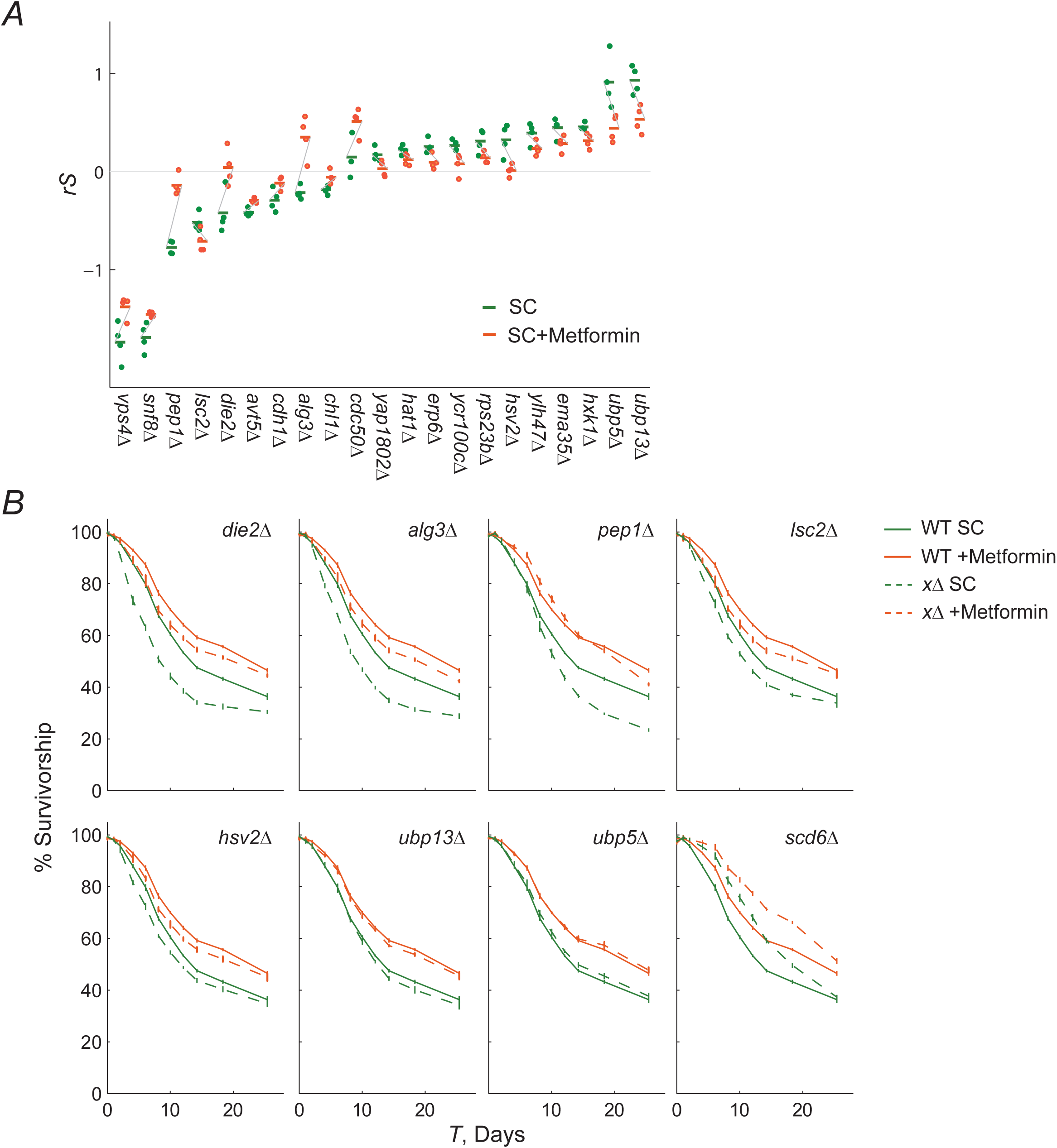
Candidate gene-drug interactions and validation. ***A*,** Plot shows individual and average measurements of *rS* of strains with differential phenotypic effects (*p*<0.05, *t*-test). Gray diagonal lines are used to visualize the sign of the gene-drug interaction. ***B*,** Validation of gene-drug interactions using the OD kinetics method. Survival curves were determined through aging monocultures of eight gene-deletion strains with some of the most extreme short- or long-lived phenotypes in SC, including candidate gene-drug interactions. Average percentage of survivorship is shown for the wild-type (solid lines) and gene-deletion strains (dashed lines) in both conditions, SC (green) and SC+Metformin (orange); error bars indicate the S.E.M (*n*=4).

